# Domino-like propagation of collective U-turns in fish schools

**DOI:** 10.1101/138628

**Authors:** Valentin Lecheval, Li Jiang, Pierre Tichit, Clément Sire, Charlotte K. Hemelrijk, Guy Theraulaz

**Affiliations:** Centre de Recherches sur la Cognition Animale, Centre de Biologie Intégrative (CBI), Centre National de la Recherche Scientifique (CNRS) & Université de Toulouse (UPS), F-31062 Toulouse, France; Groningen Institute for Evolutionary Life Sciences, University of Groningen, Centre for Life Sciences, Nijenborgh 7, 9747AG Groningen, The Netherlands; School of Systems Science, Beijing Normal University, Beijing, 100875, China; Laboratoire de Physique Théorique, CNRS & Université de Toulouse (UPS), 31062, Toulouse, France

## Abstract

Moving animal groups such as schools of fish or flocks of birds often undergo sudden collective changes of their travelling direction as a consequence of stochastic fluctuations in heading of the individuals. However, the mechanisms by which these behavioural fluctuations arise at the individual level and propagate within a group are still unclear. In the present study, we combine an experimental and theoretical approach to investigate spontaneous collective U-turns in groups of rummy-nose tetra (*Hemigrammus rhodostomus*) swimming in a ring-shaped tank. U-turns imply that fish switch their heading between the clockwise and anticlockwise direction. We reconstruct trajectories of individuals moving alone and in groups of different sizes. We show that the group decreases its swimming speed before a collective U-turn. This is in agreement with previous theoretical predictions showing that speed decrease facilitates an amplification of fluctuations in heading in the group, which can trigger U-turns. These collective U-turns are mostly initiated by individuals at the front of the group. Once an individual has initiated a U-turn, the new direction propagates through the group from front to back without amplification or dampening, resembling the dynamics of falling dominoes. The mean time between collective U-turns sharply increases as the size of the group increases. We develop an Ising spin model integrating anisotropic and asymmetrical interactions between fish and their tendency to follow the majority of their neighbours nonlinearly (social conformity). The model quantitatively reproduces key features of the dynamics and the frequency of collective U-turns observed in experiments.

## 1 Introduction

The flexible coordination of fish in schools brings important benefits [1–3]. A striking consequence of this flexibility is the performance of rapid and coherent changes in direction of travel of schools, for instance as a reaction to a predator in the neighbourhood [4, 5]. In many species, it is only a small number of individuals that detects the danger and changes direction and speed, initiating an escape wave that propagates across the entire school [6, 7]. Besides, sudden collective changes of the state of a school may also happen without external cause as a consequence of stochastic effects [8]. In these cases, local behavioural changes of a single individual can lead to large transitions between collective states of the school, such as between the schooling and milling states. Determining under what conditions fluctuations in individual behaviour, for instance in heading direction, emerge and propagate within a group is a key to understanding transitions between collective states in fish schools and in animal groups in general.

Only few theoretical and experimental studies have addressed these questions [9, 10]. Calovi et al. [10] used a data-driven model incorporating fluctuations of individual behaviour and attraction and alignment interactions between fish to investigate the response of a school to local perturbations (i.e. by an individual whose attraction and alignment behaviour differs from that of the rest of the group). They found that the responsiveness of a school is maximum near the transition region between the milling and schooling states, where the fluctuations of the polarisation are also maximal. This is entirely consistent with what happens in inert physical systems near a continuous phase transition. For instance, in magnetic systems, the polarisation of the atomic spins of a magnet near the transition point has diverging fluctuations and response to a perturbation by a magnetic field. The fluctuations of school polarisation are also expected to be strongly amplified at the transition from schooling to swarming observed when the swimming speed of individuals decreases [11]. During such a transition, the behavioural changes of a single individual are more likely to affect the collective dynamics of the school. However, the tendency of fish to conform to the speed and direction of motion of the group can also decrease the fluctuations at the level of the group with increasing group size [12]. Social conformity refers to the nonlinear response of individuals to adjust their behaviour to that of the majority [13–15].

In the present work, we analyse in groups of different size under which conditions individual U-turns occur, propagate through the group, and ultimately lead to collective U-turns. We let groups of rummy-nose tetra (*Hemigrammus rhodostomus*) swim freely in a ring-shaped tank. In this setup, fish schools can only head in two directions, clockwise or anticlockwise, and they regularly switch from one to the other. In a detailed analysis of empirical data, we reconstruct individual trajectories of fish and investigate the effect of group size on both the tendency of individuals to initiate U-turns and the collective dynamics of the U-turns. We develop an Ising-type spin model, a simple model for magnets in the physical context, to investigate the consequences on the dynamics and the propagation of information during U-turns, of the local conformity in heading, of the fish anisotropic perception of their environment, and of the asymmetric interactions between fish. We use tools and quantitative indicators from statistical physics to analyse the model. In particular, we introduce the notion of local (respectively, global/total) pseudo energy which, in the context of fish school, becomes a quantitative measure of the “discomfort” of an individual (respectively, of the group) with respect to the swimming direction of the other fish.

## 2 Material and Methods

### 2.1 Experimental procedures and data collection

A group of 70 rummy-nose tetras (*Hemigrammus rhodostomus*) were used in our experiments. This tropical freshwater species swims in a highly synchronised and polarised manner. Inside an experimental tank, a ring-shaped corridor 10 cm wide with a circular outer wall of radius 35 cm was filled with 7 cm of water of controlled quality (Electronic Supplementary Material, Figure S1A). For each trial, *n* fish were randomly sampled from their breeding tank (*n* ∈ {1, 2, 4, 5, 8, 10, 20}). Each fish only participated in a single experiment per day. For each group size, we performed between 9 and 14 replications (see Electronic Supplementary Material, Table S1). Trajectories of the fish were recorded by a Sony Handy-Cam HD camera filming from above the set-up at 50Hz in HDTV resolution (1920*×*1080p). Finally, we tracked the positions of each individual using idTracker 2.1 [16], except for groups of 20 fish. Details about experimental set-up, data extraction, and pre-processing are given in Electronic Supplementary Material.

### 2.2 Detection and quantification of individual and collective U-turns

Since fish swim in a ring-shaped tank, their heading can be converted into a binary value: clockwise or anticlockwise. Before a collective U-turn, the fish are all moving in the same direction, clockwise or anticlockwise. When one fish changes its heading to the opposite direction, it can trigger a collective U-turn (Electronic Supplementary Material, Movie S1).

From the heading angle *ϕ*_*i*_(*t*) and angular position *θ_i_*(*t*) of an individual *i* at time *t* (Electronic Supplementary Material, Figure S2), the angle of the fish relative to the wall is computed as

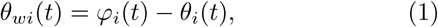

and thus the degree of alignment to the circular wall can be defined as

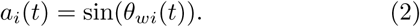

The degree of alignment *a*_*i*_(*t*) between a fish *i* and the outer wall is 1 when it is moving anticlockwise, *−*1 when moving clockwise and 0 when it is perpendicular to the wall. When a group of fish makes a collective U-turn, the degree of alignment to the wall averaged over all individuals of the group ā(*t*) changes sign. We used this as the criterion for detecting collective U-turns automatically from the smoothed time-series of ā(*t*) using a centred moving average over 9 consecutive frames. Figure 1A shows individual trajectories during a typical collective U-turn in a group of 4 fish and Figure 1B reports the corresponding evolution of the degrees of alignment *a*_*i*_(*t*). Further details about U-turns detection and the calculation of the quantities of interest are detailed in Electronic Supplementary Material, Methods.

**Figure 1.**
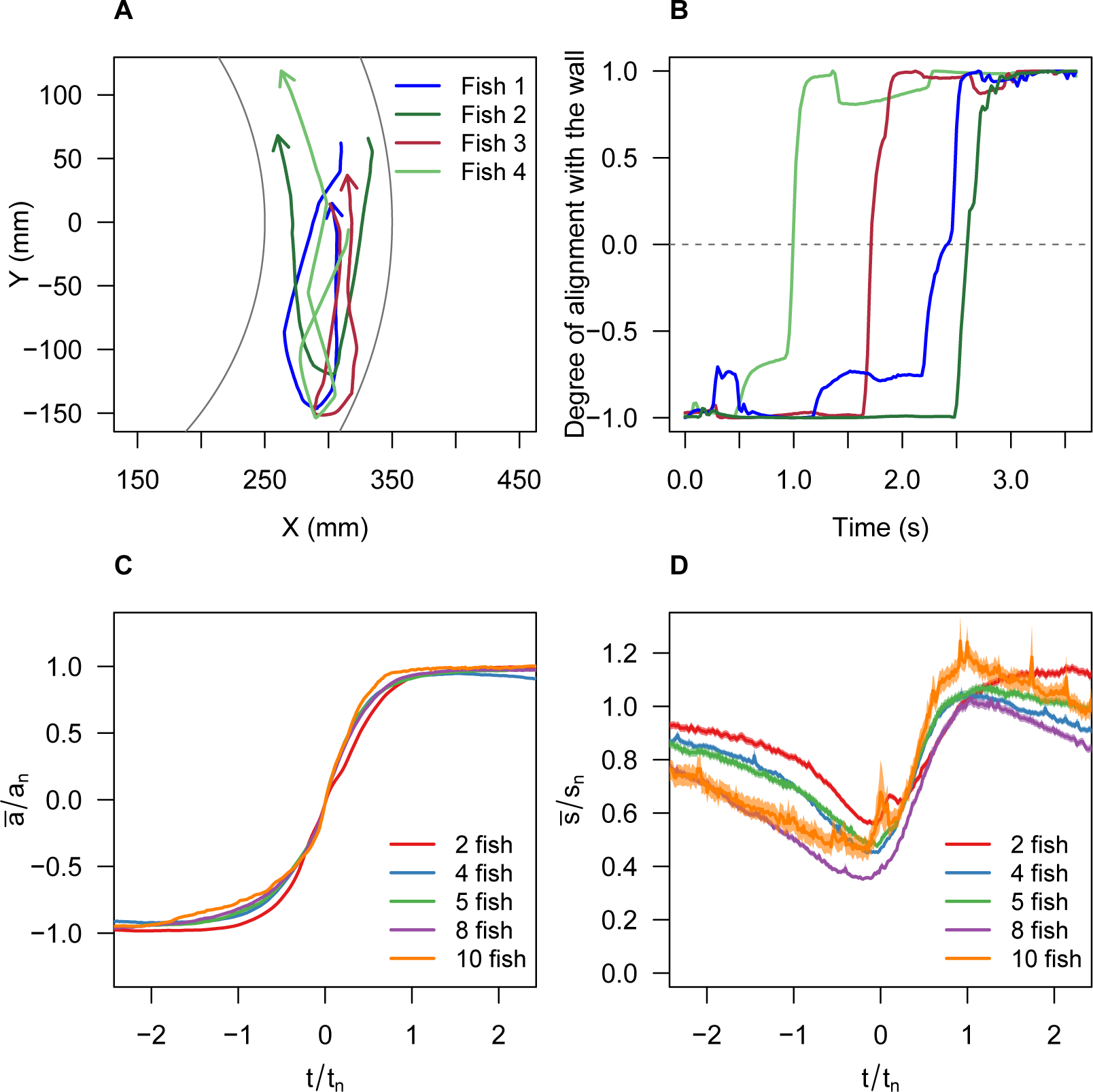
Individual trajectories (A) and degree of alignment *a*_*i*_(*t*) of fish with the wall (B) during a U-turn in a group of 4 fish. C) Normalised degree of alignment with the wall, averaged over all fish and U-turns, against the rescaled time *t/t_n_* for groups of size *n*, where *t*_*n*_ is a measure of the mean duration of a U-turn. *t* = 0 is set when *ā/a*_*n*_ = 0. D) Average individual speed 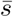 normalised by the average speed *S*_*n*_of the group, against the rescaled time *t/t_n_.*

## 3 Results

### 3.1 Spatiotemporal dynamics of collective U-turn

*Hemigrammus rhodostomus* fish form highly cohesive schools during our experiments (Electronic Supplementary Material, Figure S4A) and adjust their speed and heading to that of their group members. In a former study [17], we have shown that this is achieved through attraction and alignment interactions that have been precisely measured. Figure 2 indicates that the average time interval between two U-turns in groups of 10 fish (one U-turn every 20 min) is two orders of magnitude larger that in groups of 2 fish (one U-turn every 0.2 min). In experiments in which no collective U-turn was observed (grey triangles on Figure 2), we took the total period of observation as the interval until the next U-turn. Therefore, the average time *λ*_*n*_ between U-turns measured in groups of 4, 8, 10, and 20 fish are slightly underestimated. Thus, as group size increases, the number of collective U-turns decreases, possibly because the propensity of a fish to initiate or propagate a U-turn decreases. Like in many other species, individual fish tend to adopt the behaviour of the majority of the group members and thus inhibit the initiation of U-turns [12].

**Figure 2.**
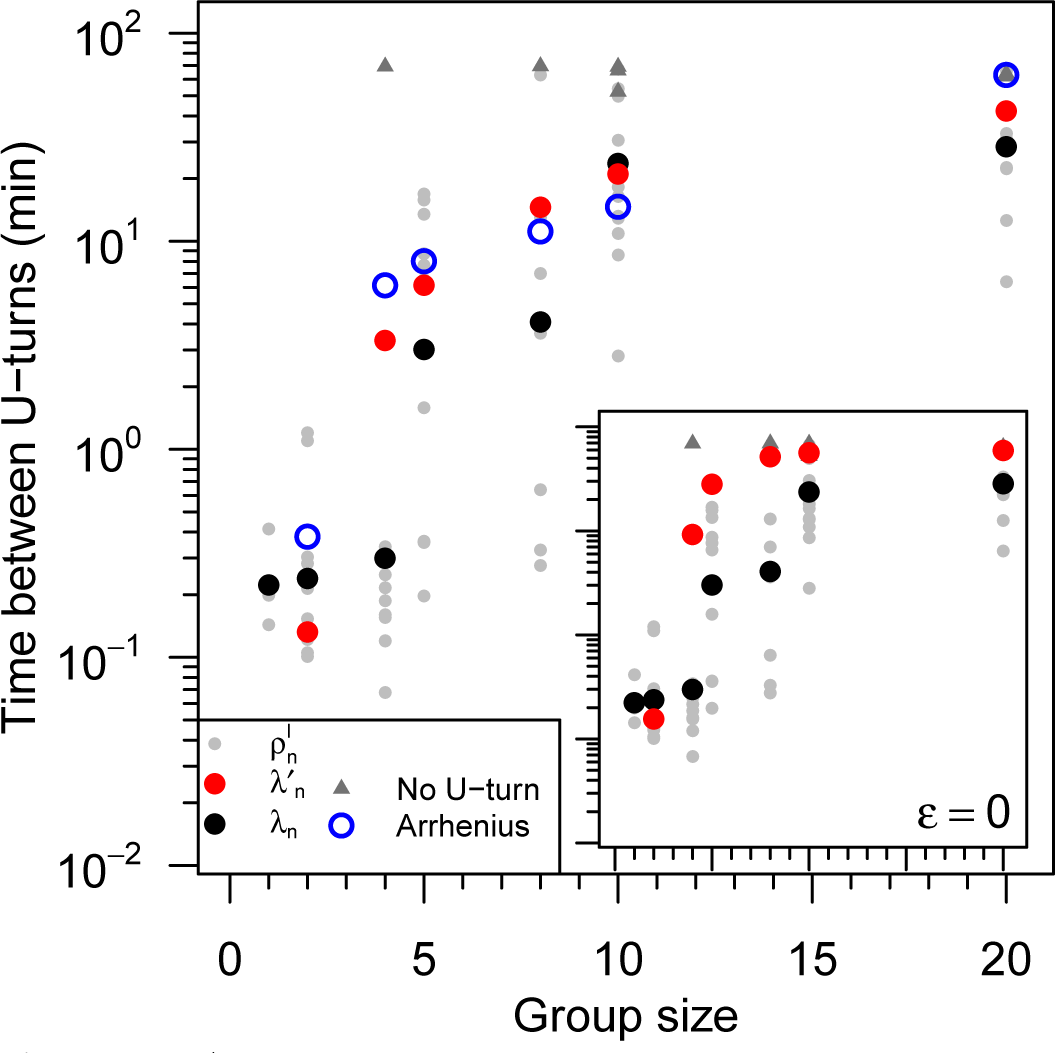
Average time between two consecutive collective U-turns as a function of group size. Average time between collective U-turns 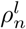 in each experiment *l* with *n* fish defined as the duration of an experiment 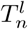 divided by the number of collective U-turns performed during this experiment (grey dots). Experiments without any collective U-turn are indicated by grey triangles, with 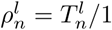. Average of the log of the time between collective U-turns over all experiments 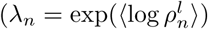; black dots) and over 1000 simulations (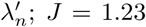 and ∈ = 0.31; red dots). Prediction of the Arrhenius law (open blue circles). Inset: results of the model without asymmetric influence (J = 1.23 and ∊ = 0).

As shown in Figure 1C, the dynamics of collective U-turns, and in particular the evolution of the mean alignment 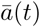, is similar for all group sizes, once time is rescaled by the mean U-turn duration (see Electronic Supplementary Material for the method used to compute the scaling parameter *t*_*n*_, which is an effective measure of the U-turn duration). In Electronic Supplementary Material, Figure S5 shows that *t*_*n*_ increases approximately linearly with group size *n.* In groups of all sizes, fish progressively decrease their velocity before turning collectively and accelerating sharply (Figure 1D). The duration of this deceleration (and then acceleration) phase is much longer than the time for the group to complete a U-turn (compare Figure 1C and Figure 1D). Moreover, the speed minimum of the group in Figure 1D is reached near the midpoint of the U-turn, when *t* = 0 and the mean alignment is *ā* = 0 in Figure 1C.

Our results show that the propagation of information is on average sequential, both in space and time. This resembles a chain of falling dominoes, for which the time interval between successive falls is constant, without any positive feedback.

Collective U-turns are usually initiated at the front of the school and the change of swimming direction propagates towards the rear (Figures 3A and B and Electronic Supplementary Material, Figure S7 and Table S2 for statistical tests). Note that Figure 3B does not show the actual shape of groups but only the average and relative positions of fish. In particular, the x-coordinates are not perfectly centred on 0 (the centroid of the average positions) for all turning ranks because the foremost fish tends to swim significantly closer to the outer wall than the fish swimming at the rear, in line with previous results in groups of two fish in the same species [17] (Electronic Supplementary Material, Table S3 for statistical tests).

**Figure 3.**
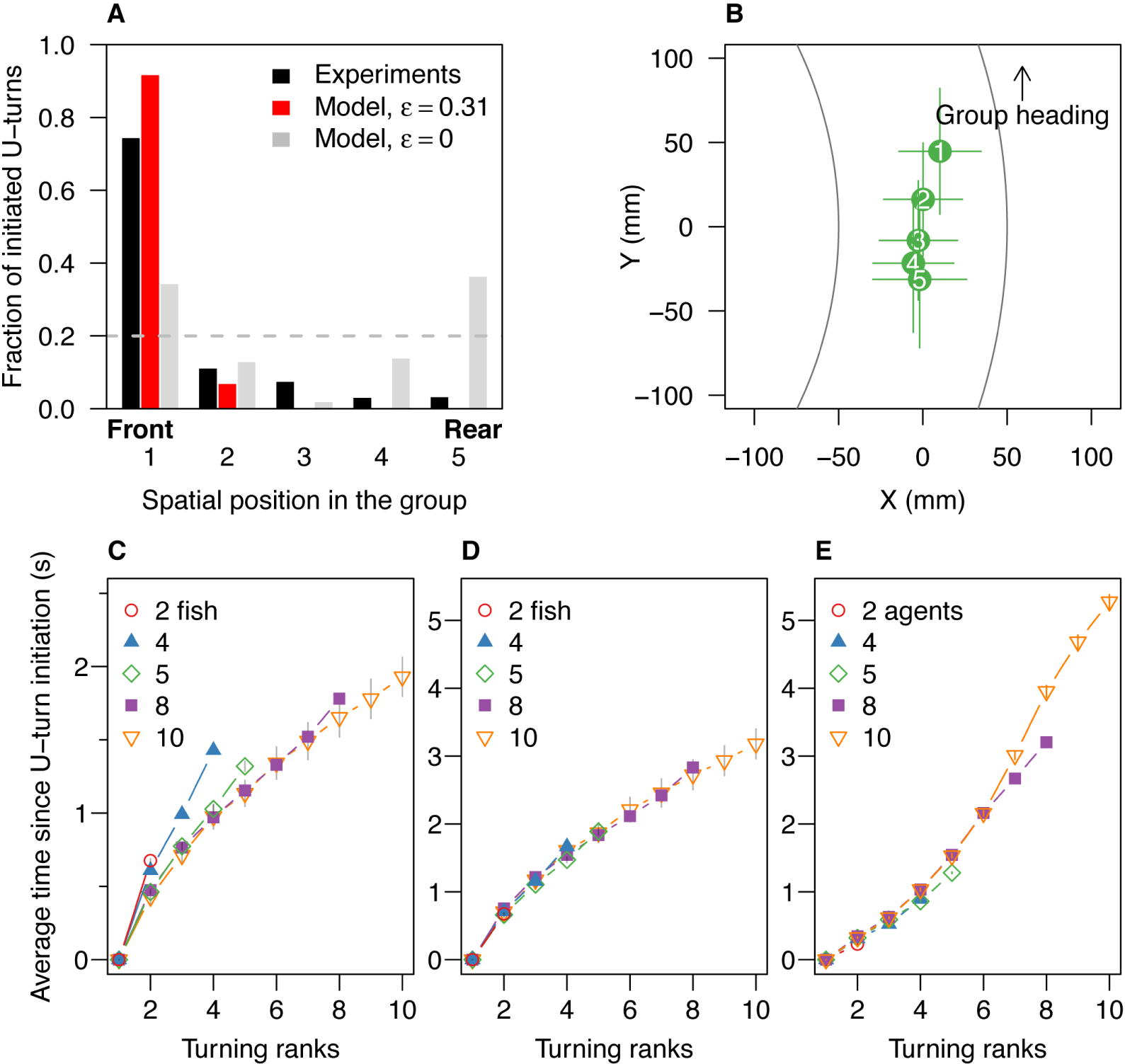
Spatiotemporal propagation of collective U-turns. A) Spatial position distribution of the initiator in groups of 5 fish in experiments (black) and in simulations with asymmetric influence (*J* = 1.23 and ∊ = 0.31; red) and without asymmetric influence (*J*= 1.23 and ∈ = 0; grey). Spatial positions go from 1 (position at the front) to 5 (position at rear). Dashed line shows the uniform distribution 1/5 = 0.2, when spatial position has no effect on the initiation of collective U-turns. B) Average relative positions(± sd) of all individuals at initiation of collective U-turns, ranked by order of turning (i.e. rank 1 is initiator) in groups of 5. Positions have been corrected so that all groups move in the same direction, with the outer wall at their right hand-side. The origin of the coordinate system is set to the centroid of the average positions of individuals. Average time interval since the beginning of a collective U-turn as a function of turning rank and group size in experiments (C and D) and in simulations (E). In D, the time is scaled by the factor *r*_*n*_ = *s*_*n*_/s_2_, where *s*_*n*_ is the average speed of groups of size *n*, revealing a behaviour almost independent of *n.*

Although the time interval between the turning of the first and the second fish is a bit longer than it is for others, the time interval between the successive turns of individuals is almost constant in a given group size (Figure 3C). This can be derived from the fact that the time since the initiation of the collective U-turn increases linearly with the turning rank. The linear propagation of information in all group sizes suggests that there is no amplification of the individual tendency to perform a U-turn: the time between two successive individuals performing U-turns does not decrease with the number of fish that already performed a U-turn. This suggests that individuals only pay attention to a small number of neighbours at a given time as was shown in [18].

The mean time interval between two successive individual U-turns decreases with group size (see Figure 3C where the slopes decrease with *n*, or Electronic Supplementary Material, Figure S9). However, when these time intervals are multiplied by a factor *r*_*n*_ proportional to the average speed *s*_*n*_ of groups of size *n (r*_*n*_ = *s*_*n*_/*s*_2_), they collapse on the same curve (Figure 3D). This suggests that the shorter reaction time of fish in larger groups is mostly due to their faster swimming speed. Larger groups swim faster (Electronic Supplementary Material, Figure S4B), presumably because fish are interacting with a greater number of neighbours and are closer to each other (Electronic Supplementary Material, Figure S4C). Indeed, fish have to avoid collisions with obstacles and other fish and the faster they swim, the shorter their reaction time, a well-known psycho-physiological principle [19].

In summary, our results show that U-turns are mostly initiated by fish located at the front of the school. U-turns are preceded by a decrease in the speed of the group. Once the U-turn has been initiated, the wave of turning propagates in a sequential way, suggesting that fish mainly copy the behaviour of a small number of individuals.

## 4 Modelling collective U-turns

### 4.1 Model description

We now introduce an Ising-type spin model [20,21] to better understand the impact of social conformity, anisotropy and asymmetry of interactions, and group size on the propagation of information during U-turns. Each agent *i* has a direction of motion *d*_*i*_ ∈ {−1, 1} with *d*_*i*_ = −1 representing swimming clockwise and *d*_*i*_ = 1 swimming anticlockwise. A U-turn performed by an agent *i* corresponds to a transition from *d*_*i*_ to −*d*_*i*_. In the model, the relative positions of individuals and the interaction network (i.e. the influential neighbours η_*i*_ of an agent *i*) are kept fixed in time (Electronic Supplementary Material, Figure S3). Agents are positioned in staggered rows (Electronic Supplementary Material, Figure S4D for experimental data supporting an oblong shape that becomes longer when the group size increases, as previously found by others, e.g. [22]) and only interact with their direct neighbours. The strength of interactions between an agent *i* and its neighbour *j* is weighted by a parameter *α*_*ij*_ that depends on the spatial position of *j* relatively to *i*. *α*_*ij*_ controls the anisotropy and asymmetry of the interactions between individuals, assuming that fish react stronger to frontal stimuli, in agreement with previous experimental results on *H. rhodostomus* [17]. We define *α*_*ij*_ = 1 + ∊ when agent *j* is in front of agent *i, α*_*ij*_ = 1 if *j* is at the side of *i*, and *α*_*ij*_ = 1 - ∊ if *j* is behind *i*, where the asymmetry coefficient ∊ ∈ [0, 1] is kept constant for all group sizes.

The propensity of an individual *i* to make a U-turn depends on the state of its neighbours η_*i*_ and on the interaction matrix *α*_*ij*_. The "discomfort" *E*_*i*_ of an agent *i* in a state *d*_*i*_ is given by

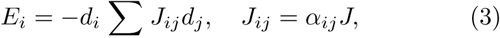

with *J*_*ij*_ the coupling constant between two neighbours *i* and *j*, set by the two positive parameters of the model, ∊ and *J* > 0. When the anisotropy of perception and asymmetry of interactions are ignored (∈ = 0), *α*_*ij*_ = 1 for all neighbouring pairs (*i, j). Ei* is minimal (and negative) when the focal fish *i* and its neighbours point in the same direction, and maximal (and positive) if the focal fish *i* points in the opposite direction of its aligned neighbours. A small value of |*E*_*i*_| corresponds to its neighbours pointing in directions nearly averaging to zero.

If an individual flips (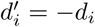), the new discomfort is 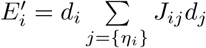, and we have

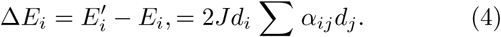

Δ*E*_*i*_ < 0 when an agent flips to the most common state of its neighbours, whereas Δ*E*_*i*_ > 0 when it flips to the state opposite to this most common state. In this ∊ = 0 case,

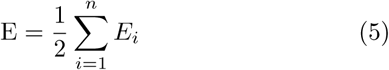

corresponds to the total actual *energy* of the magnetic system. In this context, the fully polarised state where all fish are aligned corresponds to the so-called ground-state energy, the lowest possible energy of the system. For ∊ ≠ 0, the asymmetry between the perception of *i* by *j* and that of *j* by *i* breaks this interpretation in terms of energy [17]. Yet, for ∊ *>* 0, it is still useful to define E as a pseudo energy, as will be discussed later, since it remains a good indicator of the collective discomfort of the group, i.e. the lack of heading alignment within the group.

The dynamics of the model is investigated using Monte Carlo numerical simulations inspired from the Glauber dynamics [23]. Within this algorithm, at each time step *t*_*k*+1_ = *t*_*k*_ + 1*/n* (*n* is the number of agents), an agent is drawn randomly and turns (update *d*_*i*_ to 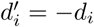) with the acceptance probability

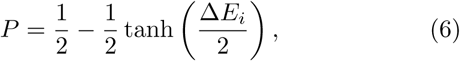

which is a sigmoid, going from *P →* 1 for Δ*Ei → −∞* (maximal acceptance if the discomfort decreases sharply), to *P →* 0 for Δ*Ei →* +*∞* (no direction switch if the discomfort would increase dramatically). The acceptance probability *P* represents the social conformity in our model and its strength (i.e. the nonlinearity of *P*) is mainly controlled by the parameter *J*(Electronic Supplementary Material, Figure S6B). For large *J >* 0, this dynamics will favour the emergence of strongly polarised states, while for *J*= 0, all fish directions will appear with the same probability during the dynamics. In physics, such a model favouring alignment between close spins is known as the Ising model, which crudely describes ferromagnetic materials, i.e. magnets.

In summary, *J* controls the directional stiffness of the fish group, while ∊ describes the fish anisotropic perception of their environment, and the asymmetric interactions between fish. After inspecting the (*J*, ∊) parameter space (see Electronic Supplementary Material, section 1.6), we find that the parameter values *J*= 1.23 and ∊ = 0.31 lead to a fair agreement between the model and experimental data, as will be shown in the next section.

### 4.2 Simulation results versus experimental data

Our model quantitatively reproduces the effect of group size on the dynamics of collective U-turns (Figure 2). This suggests that the tendency of individuals to initiate U-turns and move in the opposite direction of the whole decreases with group size. However, note the poor agreement between simulations and experimental data in groups of 4. One explanation for this may be related to the age and body size of the fish, since body size influence the strength of interactions between fish [24] (Electronic Supplementary Material, Table S1).

Even though there is no strict notion of energy in our model, we can still compute the mean pseudo energy barrier ΔE_*n*_ as a function of group size *n*. It corresponds to the mean difference between the maximum value of the pseudo energy E during the U-turn period and the reference energy computed when all the agents have the same direction (i.e. before and after the U-turn). With the interpretation of E (respectively, *E*_*i*_) as a quantitative indicator of the discomfort of the group (respectively, of the fish *i*), the (pseudo) energy barrier ΔE_*n*_ is hence a measure of the collective effort of the group to switch direction. We find that the energy barrier ΔE_*n*_ increases sublinearly with group size *n* (Electronic Supplementary Material, Figure S11). We then expect that the higher/larger is the (pseudo) energy barrier ΔE_*n*_, the more difficult it will be for the group to perform a U-turn, as it must necessarily pass through an intermediate state of greater discomfort as the group size *n* increases. As a consequence, the average time between U-turns is also expected to increase as *n* and the (pseudo) energy barrier ΔE_*n*_ increase. In fact for ∊ = 0, for which E represents a true energy, this mean time interval between direction changes is exactly given by the Arrhenius law, which can be analytically proven for our spin model evolving according to the Glauber Monte Carlo dynamics. In physical chemistry, the Arrhenius law describes for instance the switching time between two states A and B of a molecule, separated by an energy barrier associated to an intermediate state through which the molecules must necessarily pass to go from state A to state B. The Arrhenius law stipulates that the mean transition time *τ* between two states separated by an energy barrier ΔE_*n*_ grows like

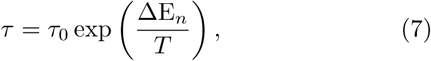

where *τ*_0_ is a time scale independent of *n* and *T* plays the role of the temperature [25]. We find that the (pseudo) Arrhenius law reproduces fairly well the experimental mean interval between U-turns as a function of group size *n*, and explains the wide range of observed time intervals (Figure 2, with *τ*_0_ *≈* 0.09 min and *T ≈* 1.9).

The sequential propagation of information is also reproduced well by the simulations of the model, both in space (Figure 3A and Electronic Supplementary Material, Figures S7 and S8) and time (Figure 3C and Electronic Supplementary Material, Figure S10). When the perception of agents is isotropic (i.e. ∊ = 0), the location of the U-turn initiation is no longer mainly at the front of the group but depends on the number of influencing neighbours (Figure 3A and Electronic Supplementary Material, Figure S3). The lower the number of influential neighbours, the higher the number of collective U-turns. For groups of 5 and ∊ = 0, the agents triggering most of the U-turns are the first and last agents because they only have two influencing neighbours.

Regarding the propagation in time, simulations reproduce the linear propagation of information at the individual scale, except for the largest group size. Figure 4A and B show that once rescaled by the U-turn duration, the average direction profile is nearly independent of the group size, and that the model prediction is in good agreement with experimental data. It takes about the same amount of time to turn the first and second half of the fish, both in experiments and in the model, although the first half of the fish is slightly slower to turn than the second half in experiments. This is consistent with the results reported on Figure 3C, where the interval between the turning of the first and the second fish was longer than between the turns of the following fish. The durations of collective U-turns are Log-normally distributed, both in experiments and in the model (Electronic Supplementary Material, Figure S12).

**Figure 4.**
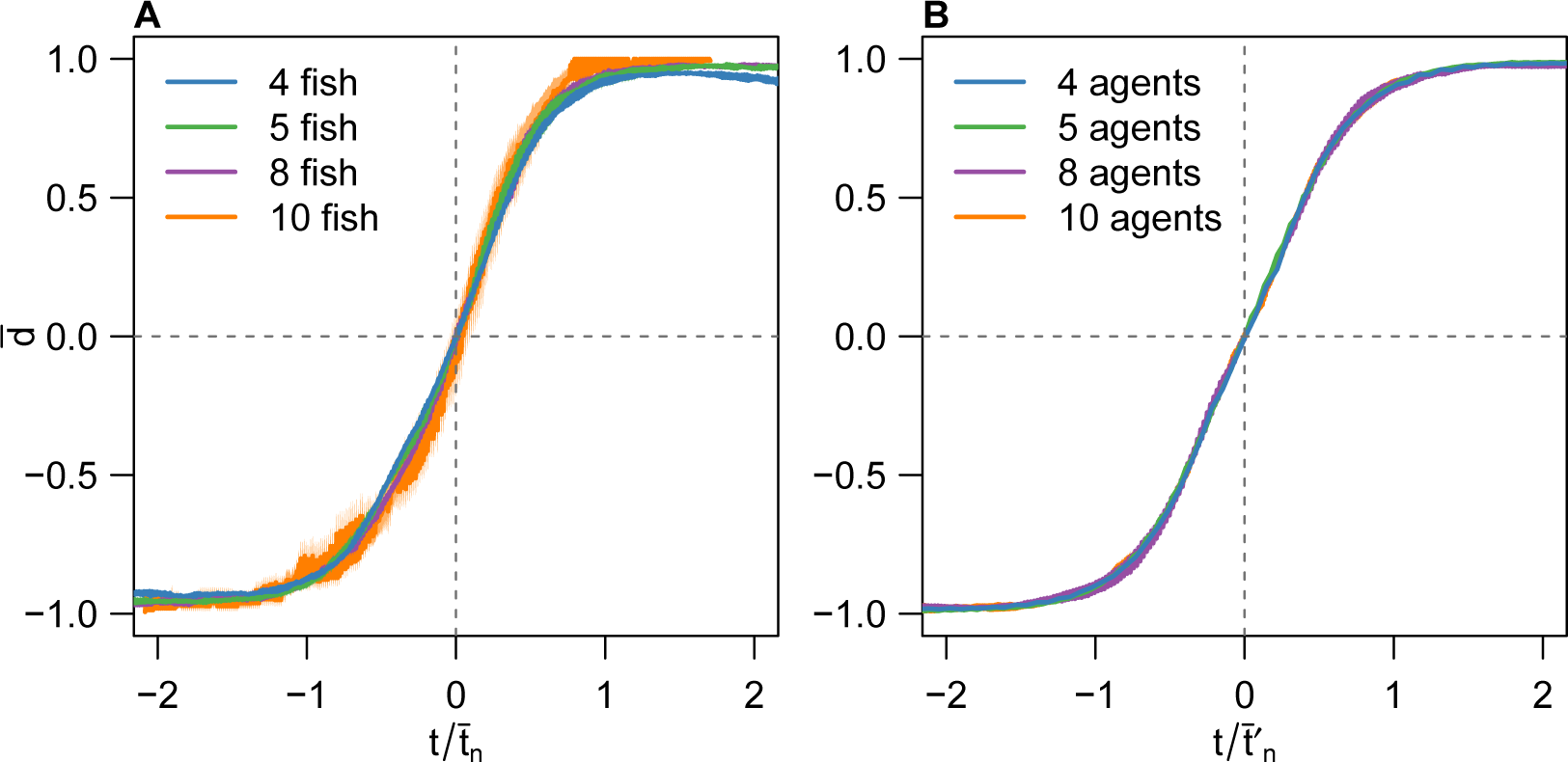
Mean swimming direction 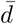 averaged over all collective U-turns as a function of scaled time 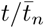 and 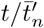 for all group sizes in (A) experiments and (B) model. *t_n_* and 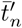 are obtained by data scaling (see Electronic Supplementary Material, Methods). The shadows stand for the standard error. In contrast to Figure 1, *t* = 0 is set to the time (*t*_*E*_ − *t*_*S*_)/2 (experiments) or 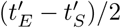 (model), where *t*_*S*_ stands for the start of the collective U-turn (first frame where at least one direction *−d*_*i*_ × *d*_0_ is positive) and *t*_*E*_ for the end of the collective U-turn (first frame where all directions *− d*_*i*_ ×*d*_0_ are positive). In A, time has also been shifted so that 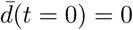.

Despite its simplicity and having only two free parameters (*J* and ∊), our model reproduces quantitatively the experimental findings, both at the collective scale (the frequency of collective U-turns, average direction profile, duration of U-turns…) and at the individual scale (the spatiotemporal features of the propagation of information). Note that a linear response of the agents to their neighbours cannot reproduce the order of magnitude of the U-turn durations measured in the experiments (Electronic Supplementary Material, Figure S6). Social conformity is thus a good candidate as an individual mechanism underlying the observed patterns including the time intervals between successive collective U-turns for different group sizes, the distribution of the U-turn duration, and the spatial propagation of information.

## 5 Discussion

How information propagates among individuals and determines behavioural cascades is crucial to understand the emergence of collective decisions and transitions between collective states in animal groups [26–29]. Here, we addressed these questions by analysing the spontaneous collective U-turns in fish schools.

We find that collective U-turns are preceded by a slowing down period. It has been shown in other fish species that speed controls alignment between individuals [11], leading slow groups to be less polarised than fast groups [8, 30–32]. In general, at slower speed, there is less inertia to turn, resulting in weaker polarisation [22, 33] and thus an increase of the fluctuations in the swimming direction of the fish [34]. Moreover, as the fish speed decreases, the fish school is in a state closer to the transition between the schooling (strong alignment) and swarming (weak alignment) states, where [32] have shown that both fluctuations in fish orientation and the sensitivity of the school to a perturbation increase. It is therefore not surprising that U-turns occur after the group has slowed down.

U-turns are mostly initiated by the fish located at the front of the group. At the front, individuals experience a lesser influence from the other fish. This is due to the perception anisotropy which results in individuals interacting more strongly with a neighbour ahead than behind. Therefore, frontal individuals are more subjects to heading fluctuations and less inhibited to initiate U-turns. Similarly, in starling flocks, the birds that initiate changes in collective travelling direction are found at the edges of the flock [35].

We found no evidence for dampening or amplification of information as fish adopt a new direction of motion. Moreover, on average, turning information propagates faster in larger groups: 0.19 s per individual in groups of 10 fish, and 0.26 s per individual in groups of 5 fish (Electronic Supplementary Material, Figure S9). This appears to be the consequence of the increase of the swimming speed with group size, which requires that individuals react faster. Indeed, our results show that the interval between successive turns of individuals during a collective U-turn decreases with swimming speed, although distance between individuals may also play a role [18]. However, the mean time interval between successive individual U-turns is almost constant and independent of the group size, once time has been rescaled by the group velocity. This points to a domino-like propagation of the new direction of motion across the group. This sequential spatiotemporal propagation of information also suggests that each fish interacts with a small number of neighbours.

We found that the level of homogeneity in the direction of motion of the schools increases with group size resulting in a lower number of collective U-turns. This phenomenon has been previously described in other fish species [12, 36] as well as in locusts in a similar set-up [37].

We developed an Ising-type spin model in which fish adopt probabilistically the direction of the majority of their neighbours, in a nonlinear way (social conformity) influenced by the anisotropic and asymmetrical interactions between fish. Since the probability that a fish chooses a direction is a nonlinear function of the number of other fish having already chosen this direction, as previously shown [38, 39], it is thus more difficult for a fish to initiate or propagate a U-turn the larger the number of fish swimming in the opposite direction [14]. The model also introduces quantitative indicators of the individual and collective discomfort (lack of alignment of heading among group members), the latter being represented by a measure of global pseudo energy of the group. Larger groups have to overcome a larger pseudo energy barrier to switch between the clockwise and anti-clockwise fully polarised states. In physics and chemistry, the fast exponential increase of the switching time between two states as a function of this energy barrier is described by the Arrhenius law, which can be actually proven using the tools of statistical physics. We find that direct numerical simulations of the model and an effective Arrhenius law both quantitatively reproduce the sharp increase of the mean time between U-turns as the group size increases. The model also shows that asymmetric interactions and the anisotropic perception of fish are not essential to explain the decrease of collective fluctuations and hence the U-turns frequency as the group size increases. Social conformity [13, 15] (controlled by the magnitude of our parameter *J*) suffices to cause fewer fluctuations with increasing group size, leading to an increased robustness of the polarised state (“protected” by increasing pseudo energy barriers).

Moreover, our model reveals that the front to back propagation of information results from the perception anisotropy and asymmetry of the fish (the *∊* parameter). Without perception anisotropy and asymmetry, U-turns are initiated by the fish that have fewer influential neighbours (in our simulations, those are the fish at the boundary of the group – all individuals would have the same probability to initiate a U-turn with periodic boundary conditions) and propagated to their neighbours without favouring any direction. Finally, the duration of a U-turn as a function of group size is quantitatively reproduced by the model, while the simulated mean direction temporal profiles during U-turns are very similar to the experimental ones, and are independent of the group size, once time is properly rescaled by the mean U-turn duration for the corresponding group size.

In summary, our work supports that social conformity, asymmetric interactions, and the anisotropic perception of fish are key to the sequential propagation of information without dampening in fish schools. Future work will be needed to disentangle the respective roles of the network topology and the actual functional forms of social interactions between fish on the propagation of information.

### Ethics statement

Experiments have been approved by the Ethics Committee for Animal Experimentation of the Toulouse Research Federation in Biology N°1 and comply with the European legislation for animal welfare.

## Contribution of authors

C.K.H. and G.T. conceived and designed the study; V.L. and P.T. performed research; V.L. and C.S. developed the model; V.L., L.J., C.S., and P.T. analysed data; V.L., C.H.K., C.S., and G.T. wrote the paper.

## Acknowledgments

We thank Daniel Calovi for his help to track experiments and his insightful comments. This study was supported by supported by the project ANR-11-IDEX-0002-02 – Transver-salité - MUSE - “Multidisciplinary study of emergence phenomena” and by grants from the CNRS and Université Paul Sabatier (project Dynabanc). V.L. was supported by doctoral fellowships from the scientific council of the Université Paul Sabatier. L.J. was funded by a grant from the China Scholarship Council (CSC NO. 201506040167).

